# The landscape of microRNA interactions annotation: analysis of three rare disorders as case study

**DOI:** 10.1101/2023.06.20.545695

**Authors:** Panni Simona, Kalpana Panneerselvam, Pablo Porras, Margaret Duesbury, Livia Perfetto, Luana Licata, Henning Hermjakob, Sandra Orchard

## Abstract

In recent years, a huge amount of data on ncRNA interactions has been described in scientific papers and databases. Although considerable effort has been made to annotate the available knowledge in public repositories, there are still significant discrepancies in how different resources capture and interpret data on ncRNAs functional and physical associations.

In the present paper, we have focused on microRNAs which regulate genes associated with rare diseases, as a case study to investigate data availability.

The list of protein-coding genes with a known role in specific rare diseases was retrieved from the Genome England PanelApp, and associated microRNA-mRNA interactions were annotated in the IntAct database, and compared with other datasets. The annotation follows recognised standard criteria approved by the IMEX consortium. RNAcentral identifiers were used for unambiguous, stable identification of ncRNAs. The information about the interaction was enhanced by a detailed description of the cell types and experimental conditions, providing a computer-interpretable summary of the published data, integrated with the huge amount of protein interactions already gathered in the database. Furthermore, for each interaction, the binding sites of the microRNA are precisely mapped on a well-defined mRNA transcript of the target gene. This information is crucial to conceive and design optimal microRNA mimics or inhibitors, to interfere in vivo with a deregulated process. As these approaches become more feasible, high-quality, reliable networks of microRNA interactions are needed to help, for instance, in the selection of the best target to be inhibited and to predict potential secondary off-target effects.

## Introduction

In recent years, it has become increasingly clear that complex and dynamic interactions between ncRNA molecules or ncRNAs and proteins contribute to virtually any biological process. MicroRNAs are probably the best characterised ncRNAs, as they can be identified with bioinformatics approaches, thanks to the conserved hairpin shape of precursor transcripts, and their possible targets can be predicted [1]. MicroRNAs regulate gene expression by guiding the RNA induced silencing complex (RISC) to those mRNAs which display sequences complementary to the microRNA “seed” i.e. nucleotides 2-7 from the 5’ end [2,3]. Predicted and verified microRNAs from more than 270 species are annotated in miRbase [4], with their precursor and mature sequences and other useful information. The latest release contains 1917 human hairpin precursors and 2654 human mature sequences, of which 26% are high-confidence microRNAs [4]. Several microRNA precursors encode two active microRNAs, named 3p and 5p respectively, which recognise different targets. Identifying the exact function for each miRNA is challenging and time consuming, and only a subset of them have been characterised. As mentioned above, it is possible to predict all possible targets for any microRNA, based on the sequence complementarity, site conservation and other features [1,5]. However, these methods provide a list of hundreds of targets for each miRNAs, many of which are not bona-fide interactors, and need to be experimentally verified [6,7]. To this aim, several techniques have been developed, for low and high throughput interaction detection, which we briefly summarise here. One of the most frequently used assays is the luciferase reporter assay, which is an adaptation of the homonymous test used to identify the regulatory regions on DNA promoters (Figure 1). The method not only permits verification of the interaction, but also the mapping of the precise binding site on the mRNA, through mutagenesis of the predicted sequence [8]. QRT-PCR and western blot allow the quantification of the mRNA and protein levels, respectively, following microRNA over-expression, and can be used to discriminate between mRNA degradation and translation inhibition, which are the two possible results of microRNA interaction (Figure 1). Pull down and immunoprecipitation approaches, commonly used to demonstrate protein binding, have been adapted to demonstrate microRNA interactions. Cross-linking ligation and sequencing of hybrids (CLASH) allows high-throughput identification of RNA-RNA interactions. In this approach, the RNA is cross-linked to the bait protein, and, after the immunoprecipitation, the ends of co-precipitated, interacting RNAs, are ligated together and sequenced to identify the couples [9]. Other high-throughput sequencing approaches of immunoprecipitated RNAs, after cross-linking to proteins of the RISC complex (CLIP-seq, PAR-CLIP etc.), provide datasets of potential RNA-RNA interactions [10], largely improving the performance of the binding predictors, although specific couples of miRNA-mRNA should be further verified [11]. While the number of publications describing microRNA interactions is constantly increasing, several databases have begun to collect them, often including high throughput indirect or weak evidence. It is worth noting that miRTarBase [12] and RAID [13,14] allow filtering of collected ncRNAs associations, to reduce false positive rates when selecting “strong evidence” interactions. The inclusion of potentially erroneous targets in network analysis may result in misleading data interpretation, as extensively discussed in Huntley et al., 2018. The UCL Functional Gene Annotation group has focused on the gene ontology annotation of human microRNAs collecting a set of highly reliable microRNA-mRNA pairs [15,16].

**Figure 1.**
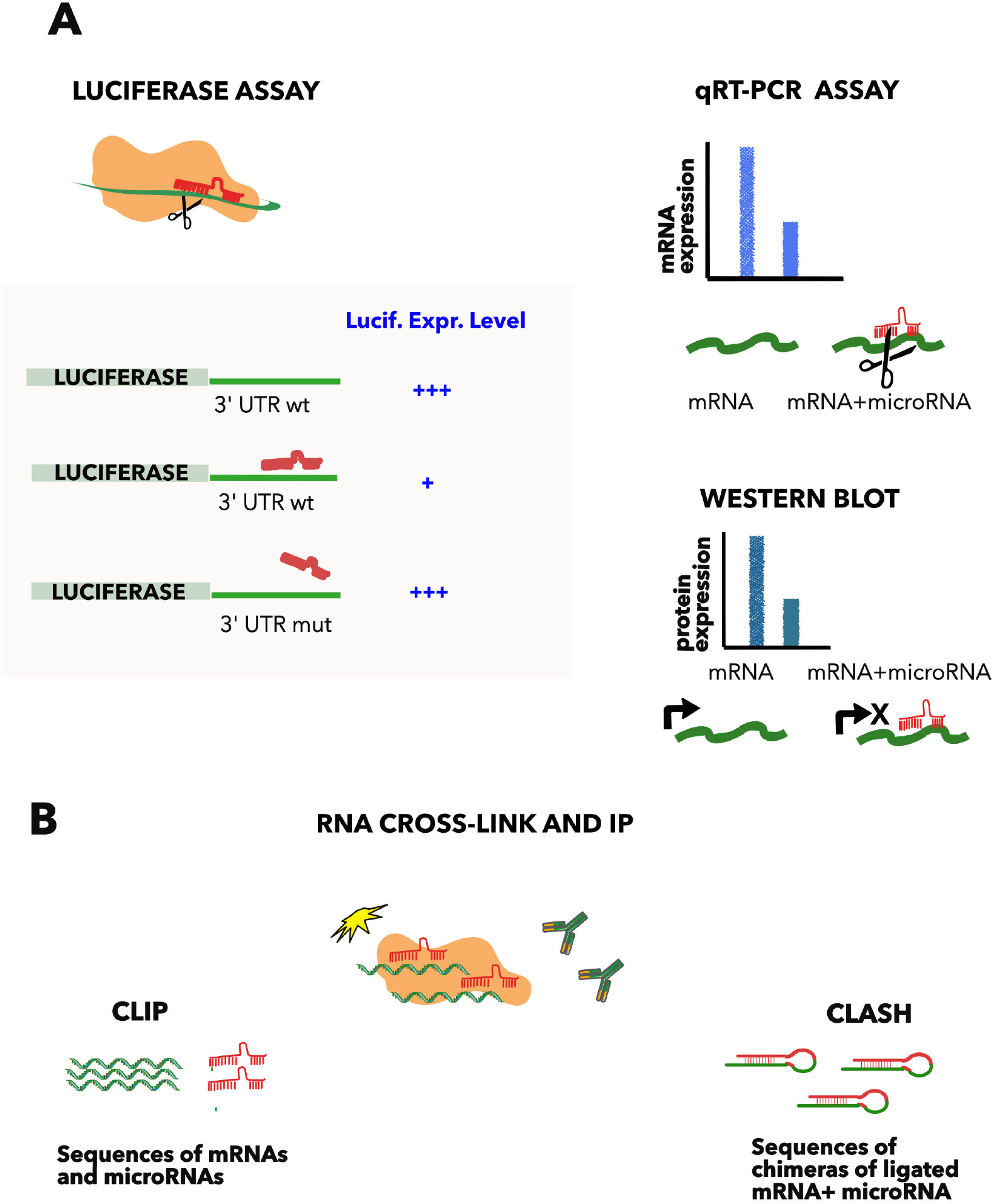
Methods commonly used to detect microRNA-mRNA interactions. A) Luciferase assay comprises of the fusion of the luciferase gene to the 3’ UTR of the microRNA target gene and the subsequent transfection of the construct with or without the microRNA. The approach demonstrates the direct interaction, compared to a mutated copy of the predicted complementary region, which acts as a negative control. Quantification of the mRNA (qRT-PCR) or protein level (WB) helps to distinguish between the mRNA degradation or the translation inhibition, but are not proof of an interaction when used in isolation. B) CLIP and CLASH. Cross-linking and immunoprecipitation of a RNA binding protein (such as AGO2), followed by RNA sequencing, allows the determination of all the RNAs bound to the protein. These approaches demonstrate direct RNA-RNA binding, if the two interacting RNAs are ligated in a hybrid before sequencing (CLASH, [9]).

Since 2002 the HUPO Proteomic Standard Initiative (HUPO-PSI) has provided a standardised annotation system for molecular interactions, and has defined the minimal information requirements and the syntax of terms used to describe an interaction experiment (MIMIx) [17], approved by members of the International Molecular Exchange (IMEx) Consortium [18,19]. Common guidelines help to elude false positives, and increase the level of detail and contextual information captured when describing molecules, such as protein binding [20]. The IntAct database, which is a member of the Consortium, has expanded its activities into the annotation of ncRNA interactions, initially focusing on *S. cerevisiae* ncRNAs [21], and subsequently on mammalian ncRNAs. In this paper, we present the collection of microRNA-mRNA interactions annotated in IntAct and we integrate and compare it with other resources. In particular, we have focused on the microRNAs that regulate genes associated with rare diseases, in order to provide information potentially useful for the development of new drugs. A rare disease is a health condition that affects a minority of people compared with other diseases that are prevalent in the population [22]. Although the increased focus on these diseases in the last few years, very little is known on microRNAs involved in rare diseases and their collection may help to gain insights into pathological mechanisms.

The interactions can be downloaded from the IntAct database (https://www.ebi.ac.uk/intact/) and also from the RNA central website (https://rnacentral.org/).

## Materials and Methods

### Data collection and deposition

MicroRNA-mRNA interactions were manually annotated in the IntAct database, according to the curation standards established by the IMEx Consortium. We retrieved relevant papers by searching the literature with text-mining seeking for the term “microRNA” (or synonymous) AND a specific gene name in the abstract, and luciferase assay or RIP, or CLASH in the methods. The collected papers were manually filtered to identify those relevant for the curation process. More than 260 papers were selected and annotated.

Genes associated with rare diseases were prioritised for annotation. The list of these genes was generated according to green panels from GenomeEngland PanelApp for the following diseases: Aniridia, Anemia Fanconi, Autism, Cakut, Cerebrak Folate Deficiency, Epidermolysis Bullosa, Familial Hirschsprung Disease, Growth Failure in Early Childhood, Early Onset Disease and Mitochondrial Disorders. These last three presented a higher number of genes regulated by microRNAs and were used in the subsequent analysis. The microRNAs are identified with RNAcentral IDs and a short label, which specify the strand of the mature microRNA [23].The database can be queried with Ensembl transcript IDs or gene common names, (then selecting mRNA) or with mRNA common names (mrna_name), as well as with the microRNA name or ID. To retrieve information on mutagenesis analysis, in order to map the binding nucleotides, from the Intact web page, click on the lens in the “Select” column of the displayed interactions, then select “Features” in the following web page. Data are also linked to RNA central [23] in the microRNA entry.

### Network analysis and databases comparison

The molecular interaction network of microRNA-mRNA interactions was downloaded from IntAct using the IntAct app [24]. To build disease-specific networks, the list of Ensembl transcript IDs was used. Networks were built and analysed with Cytoscape [25].

For comparison with other databases, miRTarBase and Raid were chosen to enable the selection of strong evidences of direct binding, comparable to those annotated in IntAct. The data were downloaded, IDs were uniformed, and the tables were uploaded in Cytoscape to build and intersect the networks.

### Functional enrichment analysis

For the “Biological Processes” enrichment analysis of microRNA targets, each transcript ID was converted to the common gene name. The tables were imported in Cytoscape to build the network, and BINGO tool [26] was used to perform the enrichment with 0.003 as significance level for GFEC and EOD related genes, while 0.05 was set for MD because there was no enrichment at lower values.

### Statistical analysis

#### Correlation analysis

A table was generated containing the number of interacting microRNAs for each gene in each of the three databases. Pearson correlation coefficient was calculated with the ggpairs function in the GGally R package [27,28]

#### Venn Diagram

Intersections among datasets were outlined by ggVennDiagram library [29] in R [28] and Jaccard index was calculated as : 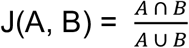 [30]

## Results

### Building microRNA network in IntAct

The main molecular function of microRNAs is to down-regulate the target gene by binding to the mRNA through few complementary nucleotides located at the 5’ of the microRNA and commonly (although not always) at the 3’ UTR of the messenger RNA. As discussed in the introduction, several experimental approaches have been developed to investigate microRNA function, but only a subset provides clear evidence of microRNA binding to mRNA, while discrimination between direct binding and causal interaction is crucial for the design of interfering drugs.

The partners of the IMEx consortium of interaction databases [18,19] have agreed to consider luciferase assay as a proof of RNA-RNA interaction, when validated with mutagenesis analysis (Figure 1). Papers describing microRNA-mRNA interactions validated with the above-mentioned methods, were selected from the literature and annotated in the IntAct database. As the interaction with mRNA results in the down-regulation of the expression, through the mRNA degradation, or in the inhibition of the translation, the microRNA is featured as “regulator” and a causality statement was used to annotate its effect.

Most of the repositories which collect microRNA interactions, identify the targets with the gene name or ID. However, microRNAs bind to messenger RNAs and, in order to map the binding sites on the target, it is necessary to refer to a specific transcript. To this aim, it is worth noting that each gene produces several messengers by alternative splicing of the precursor transcript. The number of the collected transcripts is destined to increase as new sequences will be produced from different cell-types and stages. To avoid ambiguities, we have therefore created one full-detailed entry for each target gene, cross-referred it with its Ensembl transcript ID [31], and, whenever possible, we have mapped all the annotated interactions on this entry. Figure 2 shows this process and its result. For human transcripts, among the possible transcripts listed in the Ensembl database [31] we have selected the reference shown in the GIFT curation tool (https://www.ebi.ac.uk/gifts/).

**Figure 2.**
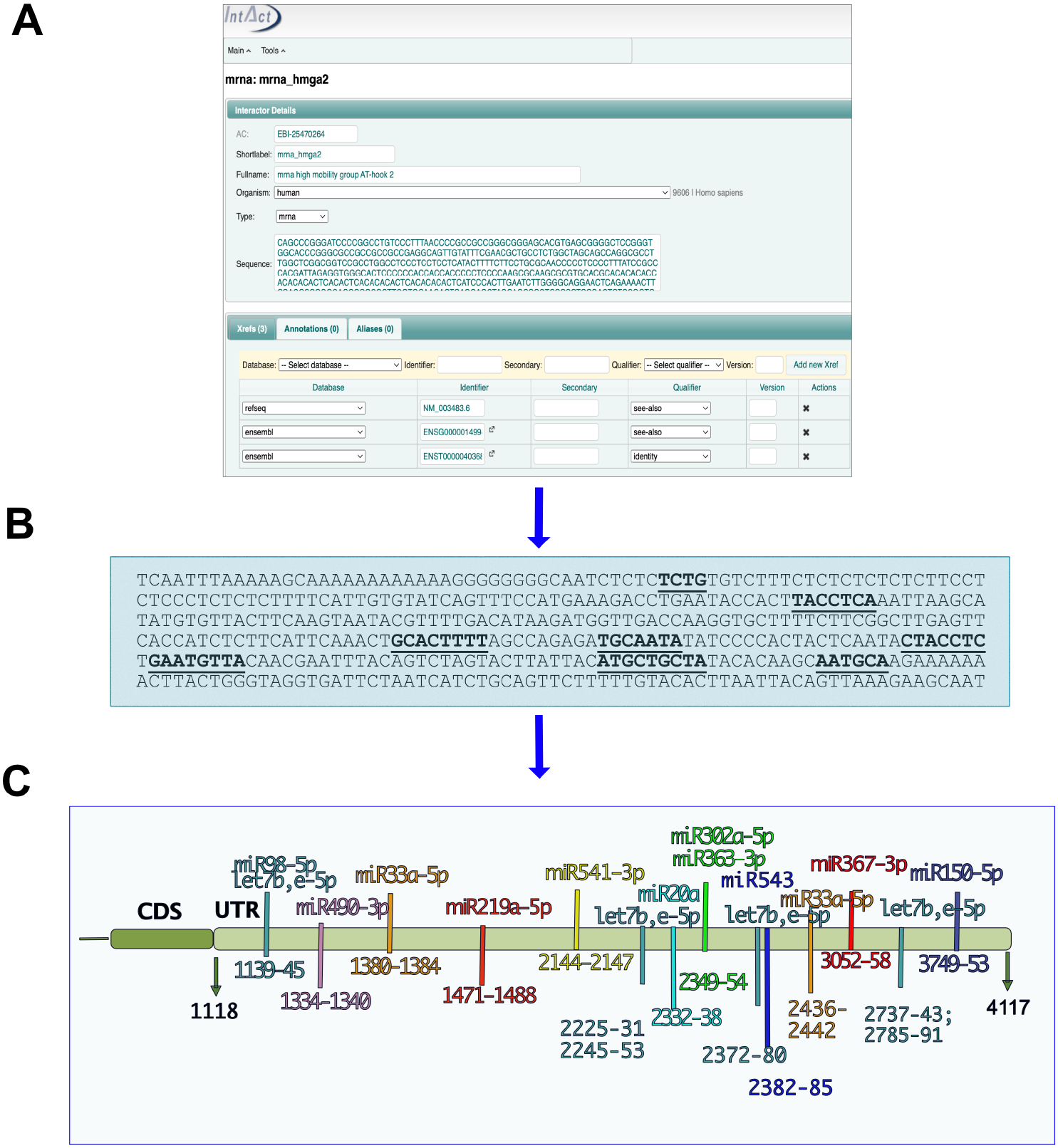
MicroRNA binding sites on mRNA_hmga2 entry. A) mRNA_hmga2 entry in IntAct is annotated with the sequence and ID of the main transcript of the gene (identified by GIFT), together with the Ensembl gene ID and a RefSeq. B) Each mutation analysis is mapped on the transcript sequence. C) Hmga2-3’UTR regions identified as necessary for the microRNA binding

We have mainly annotated human interactions, although mouse entries are also represented. Figure 3A shows a subset of the microRNA-mRNA human interaction network. Similarly to PPI networks, few nodes of both molecular types behave as hubs, presenting a number of interactions above average. The network detail blown up in Fig3B shows two connected hubs: the microRNA hsa-mir-17-5p and the mRNA_cdkn1a.

**Figure 3.**
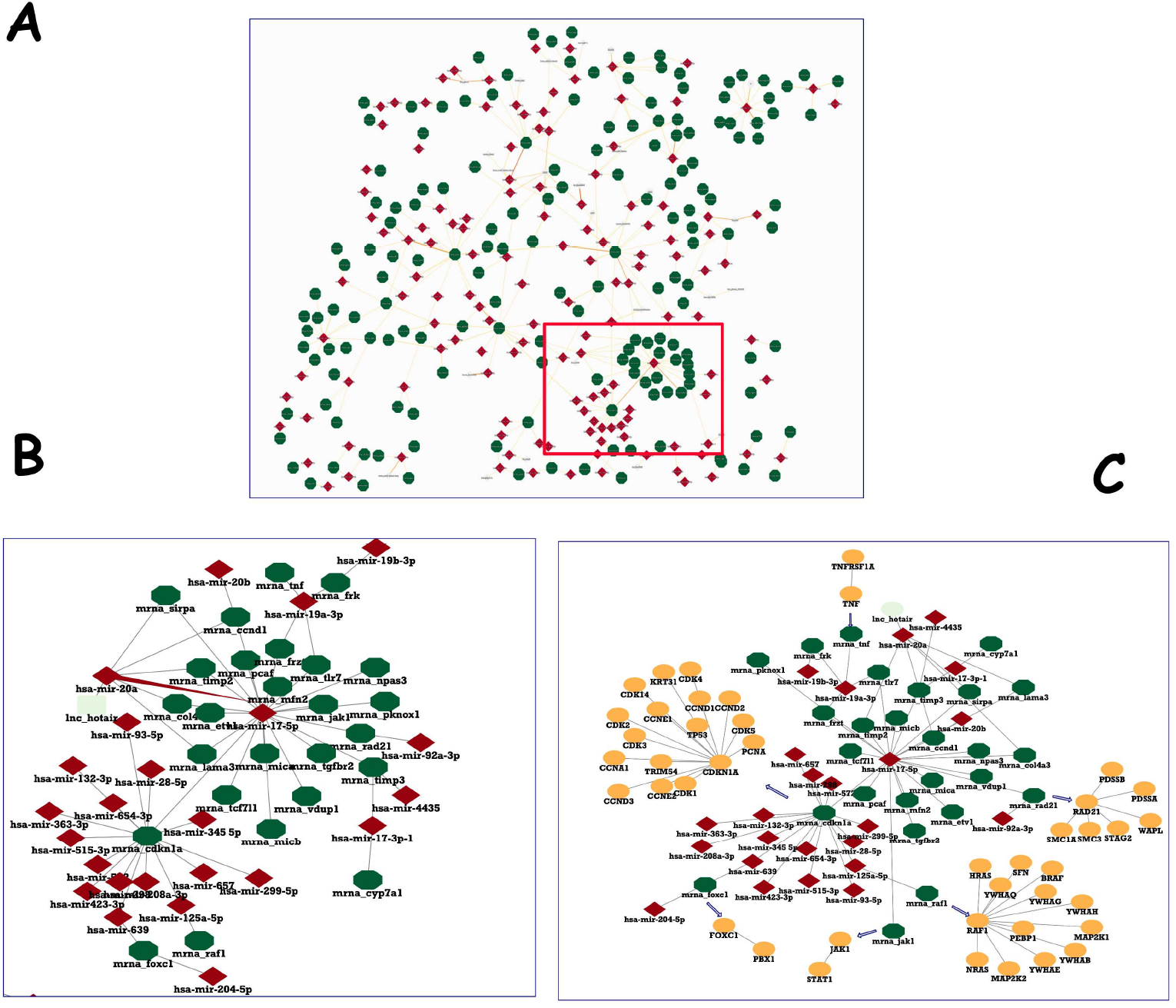
Human microRNA Network in IntAct. A) Section of the human microRNA Network, showing hubs of both entries type. B) Enlargement of a detail to show two nodes and their interactors. C) PPI network for the genes targeted by hsa-mir-17-5p was downloaded and filtered to a MIscore ⋝0.7 and merged with microRNA-mRNA interactions.

The IntAct database contains thousands of human protein interactions, each with a reliability score, so it is possible to investigate the protein complexes affected by the microRNA regulation and the connections between the protein products of co-regulated genes. Figure 3C shows some high-reliable interactors of hsa-mir-17-5p targets, that may be deregulated by mimics or inhibitors of the microRNA.

### MicroRNA regulation of rare diseases associated genes

While the amount of microRNA-interaction information is increasing continuously, very little is known about microRNAs involved in rare diseases.

Until recently, there has been limited research on the molecular mechanisms that underlie these pathologies, despite their global impact on the population and the undeniable need for optimised treatment.

We collected in IntAct a list of interactions between microRNAs and genes associated with the onset or progression of rare disorders. The list of genes was retrieved from the Genome England PanelApp [32]. Among the genes listed for each disease, few of them have been reported in the literature to bind to microRNAs. We considered several rare diseases (listed in material and methods) and, for the present study, we selected three of them, since they have a considerable number of genes regulated by microRNAs: Growth Failure in Early Childhood (GFEC), Mitochondrial Disorders (MD) and Early Onset Dementia (EOD).

Table 1 shows the list of the genes that were found to interact with microRNAs and annotated in IntAct. Growth Failure in Early Childhood is influenced by 48 genes, 17 of which were annotated in IntAct for mRNA-microRNA interactions (Figure 4). Approximately 230 genes are associated to Mitochondrial Disorders, (plus a few mitochondrial tRNA genes we did not consider). We found microRNA interactions suitable to be annotated in IntAct only for 14 of these. Finally, a list of 27 genes are associated with Early Onset Dementia, 13 of which were annotated in IntAct.

**Table 1:**
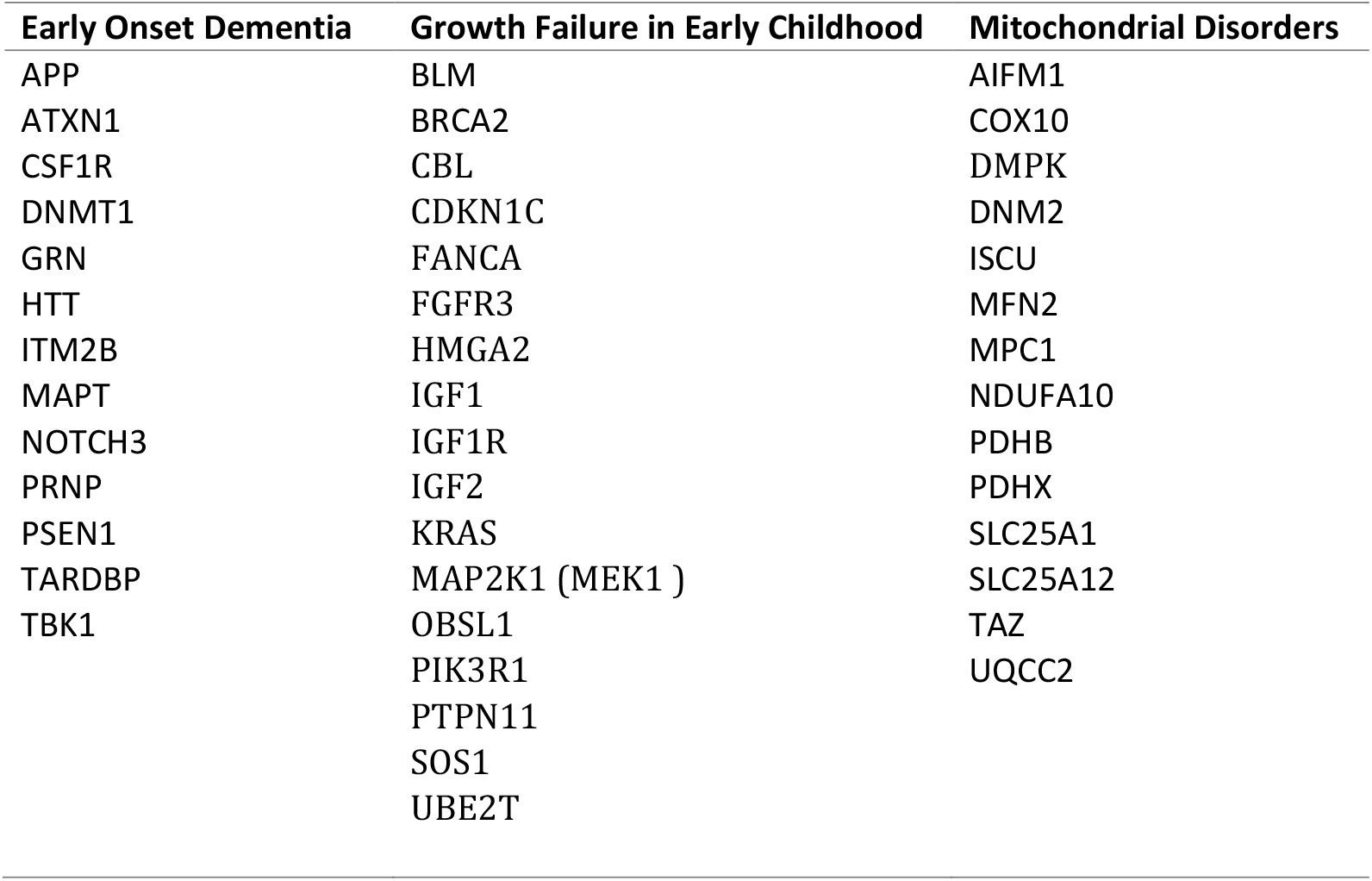
genes associated with the rare diseases and regulated by microRNAs. Only interactions confirmed by mutagenesis analysis are considered.

**Figure 4.**
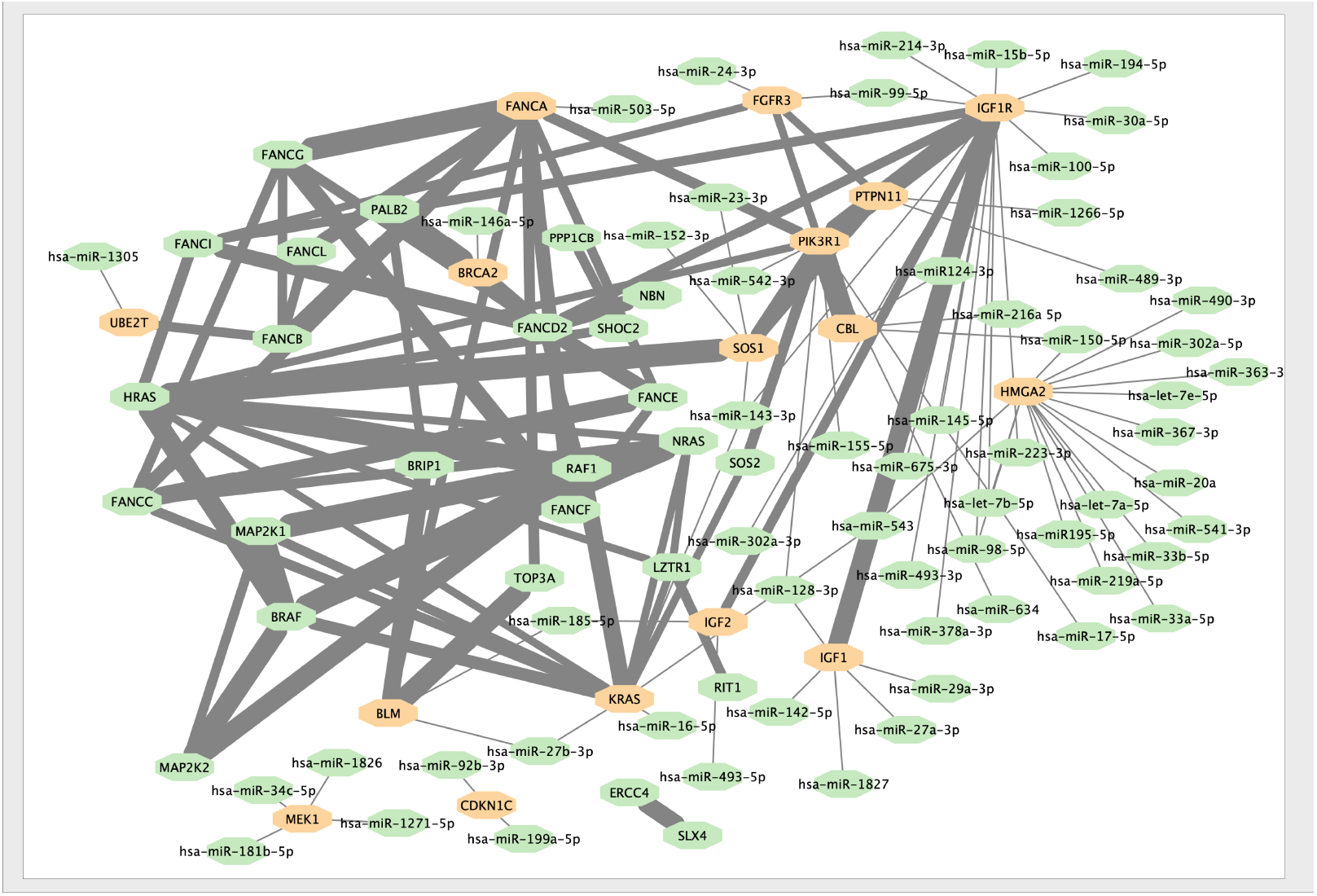
The Interaction Network of the genes associated to Growth Failure in Early Childhood. Interactions within proteins associated to GFEC merged into the microRNA-mRNA network. Proteins regulated by microRNAs are highlighted in orange. The thickness of the edge between proteins is proportional to the MI score (i.e. reliability) of the interaction [37]

We used the BINGO tool to evaluate biological processes statistically overrepresented in the microRNA regulated genes. We found an enrichment of genes involved in positive regulation of cellular processes and cell proliferation, as well as proteins involved in signalling pathways, in genes associated with GFEC and EOD, in relation to non-annotated genes.

A small number of genes associated with mitochondrial dysfunctions (MD) are regulated by microRNAs, and the enrichment analysis did not show significant enrichment for any term. Interestingly, however, there is a strong enrichment of specific metabolic and biosynthetic processes, such as: ATP synthesis coupled electron transport, carboxylic acid metabolic process, generation of precursor metabolites and tRNA metabolic processes, in genes associated with MD and not regulated by microRNAs.

We compared our results with two other databases: miRTarBase (https://mirtarbase.cuhk.edu.cn/~miRTarBase/miRTarBase_2022/php/index.php interactions confirmed by “Luciferase assay”) and RAID (https://www.rna-society.org/raid2/ interactions confirmed by “strong evidence”). The low percentage of microRNA regulated genes was confirmed in the two datasets. Supplementary Table 1 compares genes found in each of the 3 databases to interact with microRNAs. Interestingly there is a very high correlation among the number of interacting microRNAs annotated in the 3 databases for each gene, and frequently those with no interactions in one database are not annotated in the others either (Figure 5A). This suggests that, although the coverage of annotated microRNA interactions may still be low, the networks reflect what is currently known in the literature, with more than half genes of each list not affected by microRNA regulation, many involved in one or few interactions, and a small number targeted by a huge number of microRNAs. Despite the high correlation, the three datasets do not contain the same interactions: the Venn diagram shows that only 8% of them (calculated from the union of the three datasets) are present in all databases, while each dataset contains a percentage of 21%, 15% or 13% of interactions not annotated in the others (Figure 5B). This suggests that a combined effort of different resources is necessary to get the full picture and to help advance research in gene regulation (Supplementary Figure 1).

**Figure 5.**
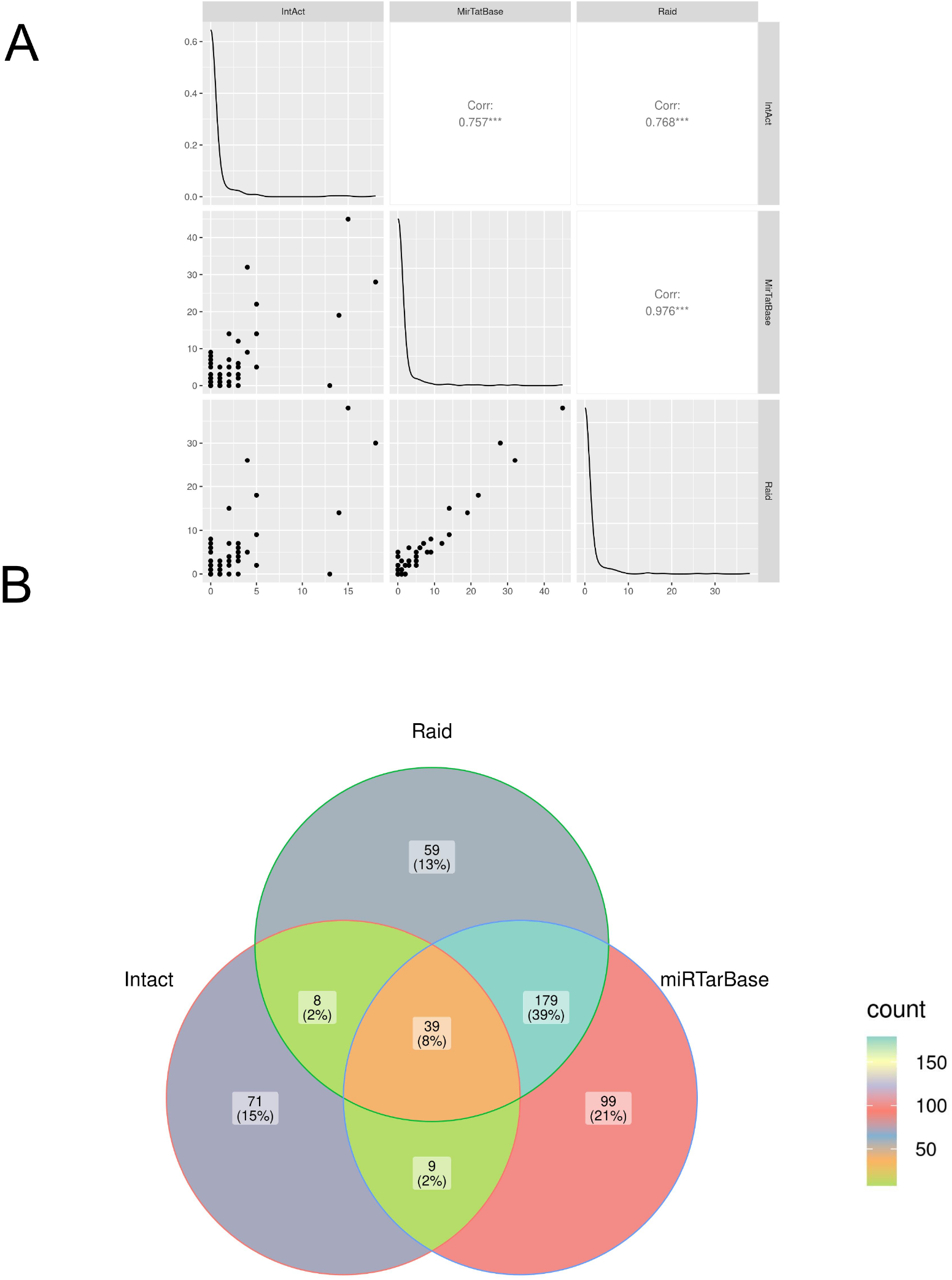
Comparison of microRNA-mRNA interactions annotated in IntAct, MirTarBase and RAID. A) Correlation analysis: the number of microRNAs regulating genes associated with the diseases in the 3 datasets was compared (see also Suppl. Table 1). B) Venn Diagram comparing interacting pairs of microRNA-mRNA annotated in the 3 datasets. The Jaccard similarity coefficient (in parenthesis) is calculated as intersection over union and indicates that only 8% of the interactions are annotated in all the datasets, and that each resource contains some exclusive information.

## Discussion

Investigations of gene-disease associations typically focus on protein variants, however, as the relevance of ncRNAs becomes more evident, their involvement in many common disorders is conceivable. For an accurate comprehension of microRNA functions, we need to assemble the entire set of miRNA-mRNA interactions, possibly annotated using standard procedures that are clearly explained to the user. Some resources collect microRNA target interactions using natural language processing to find co-occurrences of terms, followed by manual review. Despite the undeniable relevance and high coverage of these resources, there is no general acceptance of what to consider a bona-fide interactor, versus an indirect regulator. We have filtered the experimental data according to high-quality standards provided by the IMEx consortium, to collect a set of reliable microRNA targets. The IntAct database contains approximately 900 interactions involving more than 200 microRNAs. Currently modest coverage, the data set is expected to increase, as new interactions are continually being added.

The target mRNAs were annotated with the Ensembl transcript ID as identifier. In order to expedite the integration with other data identified by gene identifiers, the entries are also linked to the Ensembl gene ID. Referring to a specific transcript sequence allows the user to unequivocally annotate the interacting regions, confirmed by the mutagenesis analysis.

Recent advances in RNA molecules stabilisation and delivery methods have renewed interest in RNA-based therapies. Chemical modifications of the nucleotides increase their resistance to nucleases and encapsulation of the microRNA mimics in neutral lipids allows the delivery of the molecules into living cells or organisms [33]. In the last years several microRNA-based therapeutics have been developed, and some have entered phase II or III of clinical trials [34]. The availability of exhaustive information about microRNA binding could accelerate the design of specific mimics or inhibitors, and to predict the effect of the perturbation on the neighbourhood interactors. Since RNA-RNA interactions occur through base pairing, the design and development of interfering molecules is easier than protein inhibitors, as demonstrated by the common use of complementary molecules in scientific papers.

We have prioritised the annotation of microRNAs regulating genes involved in rare diseases, hopefully contributing to fill the gap of knowledge in this field. It is worth mentioning that the IntAct database has already collected a dataset of protein-protein interactions directly involved in rare-diseases [35]. Some diseases are directly associated with genetic mutations or variants: the disease is a direct consequence of the mutation, or, more frequently, the mutation increases the probability of the onset or progression. When the mutation directly affects a PPI interaction, the information can be retrieved from the database [36]. Very few papers describe the role of microRNA in rare diseases, and hopefully the list of interactions that affect genes associated to the diseases may help in elucidating their role. Although “rare” the “rare diseases” affect more than 30 million people in Europe, and are considered one of the major public health issues. As the return of investment in research on each individual disease may be limited, no treatment and diagnostic tests have been developed for many of them. The annotation of genes involved in rare diseases has been recognised as a crucial step to expedite the diagnosis and to establish the appropriate treatment [32]. We have expanded this knowledge by collecting existing data on the regulation of genes by microRNAs. The selected interactions are evaluated by IntAct curators and annotated according to approved CV terms. When more evidence supports an edge, a score of reliability is generated which helps the user to interpret the network [37]. The collection of “true” interactors will simplify procedures to assess the value and potential unrelated effects of a therapeutic product, and will help the design of competitors.

## Key Points

- Rare diseases are considered one of the main public health problems
- Little is known about the involvement of microRNAs in rare diseases
- MicroRNAs exert their function by regulating target genes
- We propose a method to identify microRNAs potentially involved in rare diseases
- We have collected the interactions in the IntAct database and compared them with other collections

## Funding

This work was supported by European Molecular Biology Laboratory Core Funding, National Human Genome Research Institute (NHGRI), Office of Director (OD/DPCPSI/ODSS), National Institute of Allergy and Infectious Diseases (NIAID), National Institute on Aging (NIA), National Institute of General Medical Sciences (NIGMS), National Institute of Diabetes and Digestive and Kidney Diseases (NIDDK), National Eye Institute (NEI), National Cancer Institute (NCI), National Heart, Lung, and Blood Institute (NHLBI) of the National Institutes of Health under Award Number [U24HG007822] (the content is solely the responsibility of the authors and does not necessarily represent the official views of the National Institutes of Health), Open Targets; European Union’s Horizon 2020 research and innovation program 825575 (EJP RD).

## Data availability statement

The data underlying this article are available in the IntAct database (https://www.ebi.ac.uk/intact/)

## Supplementary data Legends

**Supplementary figure 1**. Interactions between microRNAs and disease-associated genes. GFEC: Growth Failure in Early Childhood MD: Motochondrial Disorders EOD: Early Onset Dementia. Integration of the interactions from the 3 datasets are shown on the left side of the picture, Intersection on the right.

**Supplementary Table 1** Number of microRNAs interacting with the listed genes in each of the 3 databases. For Raid database “strong evidence” interactions were considered and for miRTarBase “Luciferase assay”. Data were downloaded in November 2022.

## Notes

https://www.ebi.ac.uk/intact/home

### Competing Interest Statement

The authors have declared no competing interest.

## References

1. Riffo-Campos ÁL, Riquelme I, Brebi-Mieville P. Tools for Sequence-Based miRNA Target Prediction: What to Choose? Int J Mol Sci 2016; 17:1987

2. Schirle NT, Sheu-Gruttadauria J, MacRae IJ. Structural basis for microRNA targeting. Science 2014; 346:608–613

3. Zhu L, Jiang H, Sheong FK, et al. Understanding the core of RNA interference: The dynamic aspects of Argonaute-mediated processes. Prog Biophys Mol Biol 2017; 128:39–46

4. Kozomara A, Birgaoanu M, Griffiths-Jones S. miRBase: from microRNA sequences to function. Nucleic Acids Res 2019; 47:D155–D162

5. Kern F, Backes C, Hirsch P, et al. What’s the target: understanding two decades of in silico microRNA-target prediction. Brief Bioinform 2020; 21:1999–2010

6. Witkos TM, Koscianska E, Krzyzosiak WJ. Practical Aspects of microRNA Target Prediction. Curr Mol Med 2011; 11:93–109

7. Panni S, Lovering RC, Porras P, et al. Non-coding RNA regulatory networks. Biochim Biophys Acta Gene Regul Mech 2020; 1863:194417

8. Lewis BP, Shih I -hung, Jones-Rhoades MW, et al. Prediction of mammalian microRNA targets. Cell 2003; 115:787–798

9. Helwak A, Tollervey D. Mapping the miRNA interactome by cross-linking ligation and sequencing of hybrids (CLASH). Nat Protoc 2014; 9:711–728

10. Chi SW, Zang JB, Mele A, et al. Argonaute HITS-CLIP decodes microRNA-mRNA interaction maps. Nature 2009; 460:479–486

11. Thomson DW, Bracken CP, Goodall GJ. Experimental strategies for microRNA target identification. Nucleic Acids Res 2011; 39:6845–6853

12. Huang H-Y, Lin Y-C-D, Cui S, et al. miRTarBase update 2022: an informative resource for experimentally validated miRNA-target interactions. Nucleic Acids Res 2022; 50:D222–D230

13. Lin Y, Liu T, Cui T, et al. RNAInter in 2020: RNA interactome repository with increased coverage and annotation. Nucleic Acids Res 2020; 48:D189–D197

14. Yi Y, Zhao Y, Li C, et al. RAID v2.0: an updated resource of RNA-associated interactions across organisms. Nucleic Acids Res 2017; 45:D115–D118

15. Huntley RP, Kramarz B, Sawford T, et al. Expanding the horizons of microRNA bioinformatics. RNA 2018; 24:1005–1017

16. Saverimuttu SCC, Kramarz B, Rodríguez-López M, et al. Gene Ontology curation of the blood-brain barrier to improve the analysis of Alzheimer’s and other neurological diseases. Database (Oxford) 2021; 2021:baab067

17. Hermjakob H, Montecchi-Palazzi L, Bader G, et al. The HUPO PSI’s molecular interaction format--a community standard for the representation of protein interaction data. Nat Biotechnol 2004; 22:177–183

18. Orchard S, Kerrien S, Abbani S, et al. Protein interaction data curation: the International Molecular Exchange (IMEx) consortium. Nat Methods 2012; 9:345–350

19. Orchard S, Ammari M, Aranda B, et al. The MIntAct project—IntAct as a common curation platform for 11 molecular interaction databases. Nucl. Acids Res. 2014; 42:D358–D363

20. Porras P, Barrera E, Bridge A, et al. Towards a unified open access dataset of molecular interactions. Nat Commun 2020; 11:6144

21. Panni S, Prakash A, Bateman A, et al. The yeast noncoding RNA interaction network. RNA 2017; 23:1479–1492

22. Richter T, Nestler-Parr S, Babela R, et al. Rare Disease Terminology and Definitions-A Systematic Global Review: Report of the ISPOR Rare Disease Special Interest Group. Value Health 2015; 18:906–914

23. RNAcentral Consortium. RNAcentral 2021: secondary structure integration, improved sequence search and new member databases. Nucleic Acids Res 2021; 49:D212–D220

24. Ragueneau E, Shrivastava A, Morris JH, et al. IntAct App: a Cytoscape application for molecular interaction network visualization and analysis. Bioinformatics 2021; 37:3684–3685

25. Shannon P, Markiel A, Ozier O, et al. Cytoscape: a software environment for integrated models of biomolecular interaction networks. Genome Res 2003; 13:2498–2504

26. Maere S, Heymans K, Kuiper M. BiNGO: a Cytoscape plugin to assess overrepresentation of gene ontology categories in biological networks. Bioinformatics 2005; 21:3448–3449

27. Schloerke B, Cook D, Larmarange J, Briatte F, Marbach M, Thoen E, Elberg A, Crowley J. GGally: Extension to ‘ggplot2’_. R package version 2.1.2, (2021) <Error! Hyperlink reference not valid.>.

28. R Core Team (2023). A language and environment for statistical computing. R Foundation for Statistical Computing.

29. Gao, C. ggVennDiagram: A ‘ggplot2’ Implement of Venn Diagram. R package version 1.2.2. (2022)

30. Jaccard, P. 1901. Étude comparative de la distribution florale dans une portion des Alpes et des Jura. Bulletin de la Société Vaudoise des Sciences Naturelles.

31. Martin FJ, Amode MR, Aneja A, et al. Ensembl 2023. Nucleic Acids Res 2023; 51:D933–D941

32. Martin AR, Williams E, Foulger RE, et al. PanelApp crowdsources expert knowledge to establish consensus diagnostic gene panels. Nat Genet 2019; 51:1560–1565

33. Rupaimoole R, Slack FJ. MicroRNA therapeutics: towards a new era for the management of cancer and other diseases. Nat Rev Drug Discov 2017; 16:203–222

34. Winkle M, El-Daly SM, Fabbri M, et al. Noncoding RNA therapeutics - challenges and potential solutions. Nat Rev Drug Discov 2021; 20:629–651

35. Del Toro N, Shrivastava A, Ragueneau E, et al. The IntAct database: efficient access to fine-grained molecular interaction data. Nucleic Acids Res 2022; 50:D648–D653

36. IMEx Consortium Curators, Del-Toro N, Duesbury M, et al. Capturing variation impact on molecular interactions in the IMEx Consortium mutations data set. Nat Commun 2019; 10:10

37. Villaveces JM, Jiménez RC, Porras P, et al. Merging and scoring molecular interactions utilising existing community standards: tools, use-cases and a case study. Database (Oxford) 2015; 2015:bau131

