## Supplementary figures and images for "The landscape of microRNA interactions annotation: analysis of three rare disorders as case study"

### Supplementary Figure 1

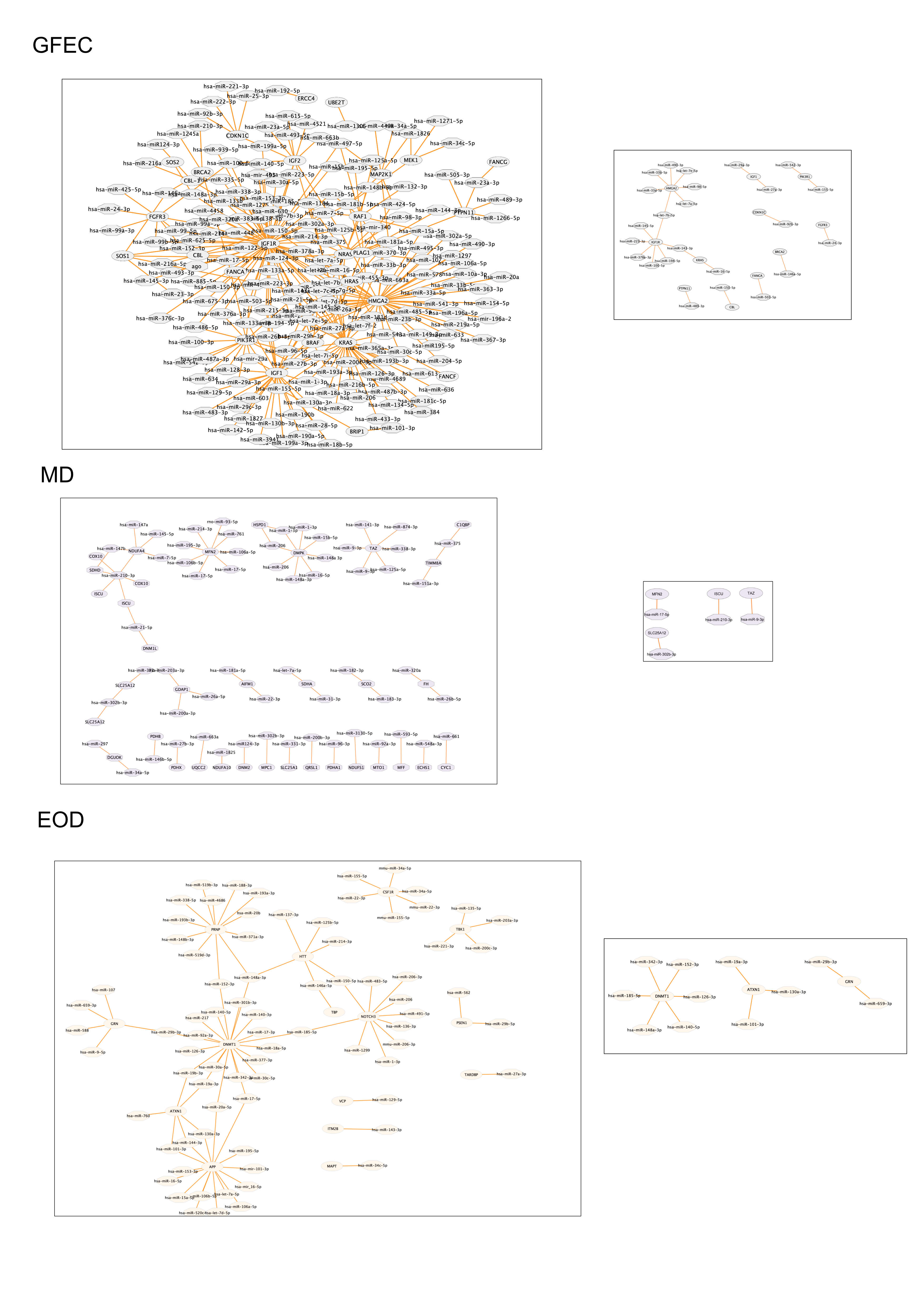
